# Genome Analysis of SARS-CoV-2 Isolate from Bangladesh

**DOI:** 10.1101/2020.05.13.094441

**Authors:** Saam Hasan, Salim Khan, Giasuddin Ahsan, Muhammad Maqsud Hossain

## Abstract

Recently the first genome sequence for a Severe acute respiratory syndrome coronavirus 2 or SARS-CoV-2 isolate from Bangladesh became available. The sequencing was carried out by the Child Health Research Foundation and provided the first insight into the genetic details of the viral strain responsible for the SARS-CoV-2 infections in Bangladesh. Here we carried out a comparative study were we explored the phylogenetic relationship between the Bangladeshi isolate with other isolates from different parts of the world. Afterwards we identified single nucleotide variants in the Bangladeshi isolate, using the Wuhan virus reference sequence. We found a total of 9 variants in the Bangladeshi isolate using 2 separate tools. Barring 2, the rest of these variants were also observed in other isolates from different countries. Most of the variants occurred in the ORF1ab gen. Another noteworthy finding was a sequence of three consecutive variants in the N protein gene that were observed in other isolates as well. Lastly the phylogenetic analysis revealed a close relationship between the Bangladeshi isolate and those from Taiwan, Kazakhstan, Greece, California, Spain, Israel, and Sri Lanka.

## Background

The Severe acute respiratory syndrome coronavirus 2 or SARS-CoV-2 has been the causative agent behind the ongoing COVID-19 pandemic. The virus which primarily infects the respiratory tract, has spread to almost every country in the world. It has infected over 4 million individuals worldwide and led to the deaths of over 283,000 [1]. A considerable body of research on the virus has already accumulated. Its genome has been sequenced in many different parts of the world. In addition, a lot of effort has already gone into identifying genetic variants. Several studies have analysed and speculated on the genetic variability of the virus and whether or not this gives it a survival advantage [2, 3, 4, 5]. Recently, the first Bangladeshi isolate of the virus has been sequenced and its data made publicly available (EPI_ISL_437912). The virus was first reported in the country last March. Since then, the number of SARS-CoV-2 infections have risen steadily, currently standing at over 17,800 infected and 269 dead [6].

As part of the ongoing situation, it becomes important for researchers to analyse and study the genome for any possible clues regarding its evolution, mutation capacity and any changes in pathogenic potential. Here, we carried out a whole genome analysis of the local viral genome sequence in order to explore its relationship with other isolates from around the world, as well as to see how it differs from them.

We tried to answer two very basic questions in this study. Firstly; what is the phylogenetic relationship between this isolate and others from around the world. This can shed light on the route taken by the virus from one country to another. As viral isolates located close to another in a phylogeny tree are more likely to share a common origin. And secondly, are there any specific genetic variants that differentiates the Bangladeshi isolate from others.

This study had two main parts. The first was the phylogeny analysis. We selected a total of 18 genomes for this, including the Bangladeshi isolate and the SARS-CoV-2 reference sequence. The rest of the genomes were chosen so as to cover as wide a geographic range as possible. We chose isolates from USA, Australia, South Korea, Japan, Israel, South Africa, Taiwan, Kazakhstan, Italy, and Spain. The second part was the variant identification. The primary goal for this component was to identify variants unique to the Bangladeshi isolate and to identify common variants that do or do not occur in the Bangladeshi isolate, so that we can understand how this strain differs from the rest.

## Methods

We used NCBI Nucleotide for obtaining the SARS-CoV-2 genomes. Search term used was “SARS-CoV-2” and the filter parameters were set to “Genomic DNA/RNA” for sequence type, and 28000-29903 base pairs for sequence length. Only complete genomes were selected.

An initial BLAST [7] run was also carried with the Bangladeshi isolate genome (EPI_ISL_437912) and the top nine hits were automatically chosen to be included in this analysis. These were isolates from USA (California, Arizona), South Africa, Spain (Barcelona), India (Hyderabad), Kazakhstan, Sri Lanka and Taiwan. Supplementary table 1 shows these hits. Along with these, we randomly selected another 8 genomes from different parts of the world. These were from India (Maharashtra), Italy, United States (Michigan), United States (Washington), Israel, Japan, South Korea and Australia. Supplementary table 2 shows the BLAST alignment summary for all these isolates, including the Bangladeshi one, with the SARS-CoV-2 RefSeq.

After selected our genome, we carried out a pairwise alignment for all of them, using the Wuhan virus reference sequence as the reference. This was done by solving the Needleman-Wunsch alignment problem for these sequences, using the Biostrings package on R [8]. A multiple sequence alignment was also carried out using the NPhylogeny.fr tool. The latter being a MAFFT program [9].

The generated alignment file from the MAFFT alignment was then used to create the phylogenetic tree for all these viral isolates. This was done using the MEGA tool for phylogenetic analysis [10]. The ML heuristic model implemented was Nearest-Neighbour-Interchange (NNI), the model used was the Tamura-Nei model, and the starting tree used was BioNJ.

For identifying SNPs, we used a combination of the MismatchTable function from Biostrings and the BasebyBase tool from the Viral Bioinformatics Research Centre [11]. We used the results from both the tools in our subsequent comparisons.

## Results

A total of 9 variants were identified in the Bangladeshi isolate by the Biostrings method and an identical number from the BasebyBase method. Both the tools gave us identical variants in terms of position and base change for the Bangladeshi isolate, lending more credibility to the variant calls. Table 1 shows the position and base changes associated with these 9 variants. 7 out of these 9 variants were also found in other isolates. As for most common variants, the C to T variant at position 241, the C to T at position 3037, and the A to G at position 23403 had the highest occurrence. All three of these were present in 12 of the 18 isolates. Other common variants included the G to T variant at position 11083 (two isolates) and a consecutive series of three variants at positions 28881, 28882, and 28883 (G to A, G to A, and G to C respectively) that were found in 6 isolates.

**Table 1:** Single Nucleotide Variants identified in the Bangladeshi SARS-CoV-2 isolate.

| Position | Bangladeshi Isolate | Reference Sequence | Gene |
| --- | --- | --- | --- |
| 241 | T | C | ORF1ab |
| 1163 | T | A | ORF1ab |
| 3037 | T | C | ORF1ab |
| 14408 | T | C | ORF1ab |
| 17019 | T | G | ORF1ab |
| 23403 | G | A | S |
| 28881 | A | G | N |
| 28882 | A | G | N |
| 28883 | C | G | N |

**Table 2:** Complete list of variants identified using Biostrings. A total of 63 unique variants were identified across the 18 isolates.. The variant at positions 241, 3037, and 23403 occurred in the most number of genomes, a total of 12.

| Isolate | Sequence Nuc | QUAL | Ref Start | Ref End | Ref Nuc | Gene |
| --- | --- | --- | --- | --- | --- | --- |
| **MT308700.1-USA-Michigan** | T | 7 | 241 | 241 | C | Intergenic |
|  | T | 7 | 1059 | 1059 | C | ORF1ab |
|  | T | 7 | 3037 | 3037 | C | ORF1ab |
|  | T | 7 | 14408 | 14408 | C | ORF1ab |
|  | G | 7 | 23403 | 23403 | A | S |
|  | T | 7 | 25563 | 25563 | G | ORF3a |
| **EPI_ISL_437912 Bangladesh** | T | 7 | 241 | 241 | C | Intergenic |
|  | T | 7 | 1163 | 1163 | A | ORF1ab |
|  | T | 7 | 3037 | 3037 | C | ORF1ab |
|  | T | 7 | 14408 | 14408 | C | ORF1ab |
|  | T | 7 | 17019 | 17019 | G | ORF1ab |
|  | G | 7 | 23403 | 23403 | A | S |
|  | A | 7 | 28881 | 28881 | G | N |
|  | A | 7 | 28882 | 28882 | G | N |
|  | C | 7 | 28883 | 28883 | G | N |
| **MT428553.1-Kazakhstan** | T | 7 | 241 | 241 | C | Intergenic |
|  | T | 7 | 3037 | 3037 | C | ORF1ab |
|  | T | 7 | 14408 | 14408 | C | ORF1ab |
|  | G | 7 | 18956 | 18956 | A | ORF1ab |
|  | T | 7 | 19170 | 19170 | C | ORF1ab |
|  | A | 7 | 19509 | 19509 | G | ORF1ab |
|  | G | 7 | 23403 | 23403 | A | S |
|  | A | 7 | 28881 | 28881 | G | N |
|  | A | 7 | 28882 | 28882 | G | N |
|  | C | 7 | 28883 | 28883 | G | N |
| **MT385418.1-USA-California** |  |  |  |  |  |  |
|  | T | 7 | 241 | 241 | C | Intergenic |
|  | T | 7 | 313 | 313 | C | Intergenic |
|  | T | 7 | 3037 | 3037 | C | ORF1ab |
|  | T | 7 | 14408 | 14408 | C | ORF1ab |
|  | G | 7 | 23403 | 23403 | A | S |
|  | T | 7 | 28086 | 28086 | G |  |
|  | A | 7 | 28881 | 28881 | G | N |
|  | A | 7 | 28882 | 28882 | G | N |
|  | C | 7 | 28883 | 28883 | G | N |
| **MT415321.1-India** |  |  |  |  |  |  |
|  | T | 7 | 241 | 241 | C | Intergenic |
|  | T | 7 | 3037 | 3037 | C | ORF1ab |
|  | T | 7 | 14408 | 14408 | C | ORF1ab |
|  | G | 7 | 23403 | 23403 | A | S |
| **MT359866.1-Spain** |  |  |  |  |  |  |
|  | T | 7 | 241 | 241 | C | Intergenic |
|  | T | 7 | 313 | 313 | C | ORF1ab |
|  | T | 7 | 3037 | 3037 | C | ORF1ab |
|  | T | 7 | 14408 | 14408 | C | ORF1ab |
|  | G | 7 | 23403 | 23403 | A | S |
|  | G | 7 | 27302 | 27302 | A |  |
|  | A | 7 | 28881 | 28881 | G | N |
|  | A | 7 | 28882 | 28882 | G | N |
|  | C | 7 | 28883 | 28883 | G | N |
| **MT324062.1-South Africa** |  |  |  |  |  |  |
|  | T | 7 | 241 | 241 | C | Intergenic |
|  | T | 7 | 3037 | 3037 | C | ORF1ab |
|  | T | 7 | 13620 | 13620 | C | ORF1ab |
|  | T | 7 | 14408 | 14408 | C | ORF1ab |
|  | T | 7 | 21595 | 21595 | C | S |
|  | G | 7 | 23403 | 23403 | A | S |
| **MT339041.1-USA-Arizona** |  |  |  |  |  |  |
|  | T | 7 | 241 | 241 | C | Intergenic |
|  | T | 7 | 1059 | 1059 | C | ORF1ab |
|  | T | 7 | 3037 | 3037 | C | ORF1ab |
|  | T | 7 | 14408 | 14408 | C | ORF1ab |
|  | G | 7 | 23403 | 23403 | A | S |
|  | T | 7 | 25563 | 25563 | G | ORF3a |
| **MT304476.1-Korea** |  |  |  |  |  |  |
|  | T | 7 | 5572 | 5572 | G | ORF1ab |
|  | T | 7 | 11083 | 11083 | G | ORF1ab |
|  | T | 7 | 14805 | 14805 | C | ORF1ab |
|  | T | 7 | 26144 | 26144 | G | ORF3a |
|  | T | 7 | 28311 | 28311 | C | N |
| **MT374116.1-Taiwan** |  |  |  |  |  |  |
|  | T | 7 | 241 | 241 | C | Intergenic |
|  | T | 7 | 3037 | 3037 | C | ORF1ab |
|  | T | 7 | 14408 | 14408 | C | ORF1ab |
|  | G | 7 | 23403 | 23403 | A | S |
|  | T | 7 | 25810 | 25810 | C | ORF3a |
|  | A | 7 | 28881 | 28881 | G | N |
|  | A | 7 | 28882 | 28882 | G | N |
|  | C | 7 | 28883 | 28883 | G | N |
| **MT328032.1-Greece** |  |  |  |  |  |  |
|  | T | 7 | 241 | 241 | C | Intergenic |
|  | T | 7 | 3037 | 3037 | C | ORF1ab |
|  | T | 7 | 14408 | 14408 | C | ORF1ab |
|  | T | 7 | 14786 | 14786 | C | ORF1ab |
|  | C | 7 | 18126 | 18126 | T | ORF1ab |
|  | G | 7 | 23403 | 23403 | A | S |
|  | A | 7 | 28881 | 28881 | G | N |
|  | A | 7 | 28882 | 28882 | G | N |
|  | C | 7 | 28883 | 28883 | G | N |
| **MT293225.1 USA-Washington** |  |  |  |  |  |  |
|  | T | 7 | 34 | 34 | A | Intergenic |
|  | T | 7 | 35 | 35 | A | Intergenic |
|  | T | 7 | 36 | 36 | C | Intergenic |
|  | H | 7 | 37 | 37 | C | Intergenic |
|  | T | 7 | 8782 | 8782 | C | ORF1ab |
|  | C | 7 | 11320 | 11320 | T | ORF1ab |
|  | T | 7 | 17747 | 17747 | C | ORF1ab |
|  | G | 7 | 17858 | 17858 | A | ORF1ab |
|  | T | 7 | 18060 | 18060 | C | ORF1ab |
|  | C | 7 | 28144 | 28144 | T | ORF8 |
| **MT066156.1-Italy** |  |  |  |  |  |  |
|  | T | 7 | 2269 | 2269 | A | ORF1ab |
|  | T | 7 | 26144 | 26144 | G | ORF3a |
| **MT050493.1 India-Maharashtra** |  |  |  |  |  |  |
|  | G | 7 | 1691 | 1691 | A | ORF1ab |
|  | T | 7 | 6501 | 6501 | C | ORF1ab |
|  | T | 7 | 8782 | 8782 | C | ORF1ab |
|  | T | 7 | 16877 | 16877 | C | ORF1ab |
|  | T | 7 | 24351 | 24351 | C | S |
|  | C | 7 | 28144 | 28144 | T | ORF8 |
| **LC528233.1-Japan** |  |  |  |  |  |  |
|  | C | 7 | 1 | 1 | A | Intergenic |
|  | C | 7 | 2 | 2 | T | Intergenic |
|  | C | 7 | 3 | 3 | T | Intergenic |
|  | C | 7 | 4 | 4 | A | Intergenic |
|  | A | 7 | 7 | 7 | G | Intergenic |
|  | T | 7 | 11083 | 11083 | G | ORF1ab |
|  | T | 7 | 29635 | 29635 | C | ORF10 |
|  | C | 7 | 29876 | 29876 | A | Intergenic |
|  | G | 7 | 29878 | 29878 | A | Intergenic |
|  | T | 7 | 29879 | 29879 | A | Intergenic |
|  | G | 7 | 29880 | 29880 | A | Intergenic |
|  | C | 7 | 29883 | 29883 | A | Intergenic |
|  | T | 7 | 29886 | 29886 | A | Intergenic |
|  | G | 7 | 29887 | 29887 | A | Intergenic |
|  | C | 7 | 29888 | 29888 | A | Intergenic |
|  | T | 7 | 29889 | 29889 | A | Intergenic |
|  | G | 7 | 29891 | 29891 | A | Intergenic |
|  | G | 7 | 29892 | 29892 | A | Intergenic |
|  | G | 7 | 29893 | 29893 | A | Intergenic |
| **MT007544.1 Australia** |  |  |  |  |  |  |
|  | C | 7 | 19065 | 19065 | T | ORF1ab |
|  | G | 7 | 22303 | 22303 | T | S |
|  | T | 7 | 26144 | 26144 | G | ORF3a |
| **MT276598.1 Israel** |  |  |  |  |  |  |
|  | T | 7 | 241 | 241 | C | Intergenic |
|  | T | 7 | 313 | 313 | C | Intergenic |
|  | T | 7 | 3037 | 3037 | C | ORF1ab |
|  | T | 7 | 14408 | 14408 | C | ORF1ab |
|  | G | 7 | 23403 | 23403 | A | S |
|  | A | 7 | 28881 | 28881 | G | N |
|  | A | 7 | 28882 | 28882 | G | N |
|  | C | 7 | 28883 | 28883 | G | N |
| **MT371048.1 Sri Lanka** |  |  |  |  |  |  |
|  | T | 7 | 241 | 241 | C | Intergenic |
|  | T | 7 | 3037 | 3037 | C | ORF1ab |
|  | T | 7 | 14408 | 14408 | C | ORF1ab |
|  | G | 7 | 23403 | 23403 | A | S |
|  | A | 7 | 28881 | 28881 | G | N |
|  | A | 7 | 28882 | 28882 | G | N |
|  | C | 7 | 28883 | 28883 | G | N |

The Bangladeshi virus showed most similarity with the viral isolate from Sri Lanka, as was previously suggested by the BLAST search. Both of these genomes contained the exact identical variants, both in terms of position and nucleotide change. Aside from that, the most interesting pattern we observed with the single nucleotide variants was the aforementioned three nucleotide change from position 28881-28883. This was located in the N protein gene of the virus. It is a gene known to play a critical role in virion assembly and structure. 8 of the isolates contained this tri-nucleotide change (Bangladesh, Kazakhstan, California, Taiwan, Greece, Spain, Sri Lanka and Israel). The other isolates did not contain variants at any of these three positions. Finally a variant at position 17019 that caused a change from T to G and a variant at position 11063 that caused a change from T to A, were only found in the Bangladeshi isolate.

A total of 63 unique variants were identified across the two runs. 41 were identified with BasebyBase and another 22 unique variants were found using Biostrings. Table 3 shows all the variants found with Biostrings. Figure 1 displays a binary heatmap of all the variants identified with BasebyBase.

As far as gene specific distribution of variants is concerned, the Bangladeshi isolate, much like all the others, contained most variants in the ORF1ab gene. This is to be expected as that is the largest gene in the SARS-CoV-2 genome, spanning over half its genome length. It also contained the previously discussed 3 variants in the N protein gene and 1 variant in the S protein gene. The latter, an A to G mutation at position 23403, has already been implicated in a number of previous studies. Previously it was believed this particular mutation is common the viral strains in Europe [12]. Although out analysis did show that isolates from other Asian countries (India, Kazakhstan, Taiwan, Sri Lanka, Israel), in addition to the Bangladeshi one, also contained this particular variant. Figure 2A and 2B show the number of variants per gene for the all the isolates overall and for the Bangladeshi isolate alone respectively.

**Figure 2:**
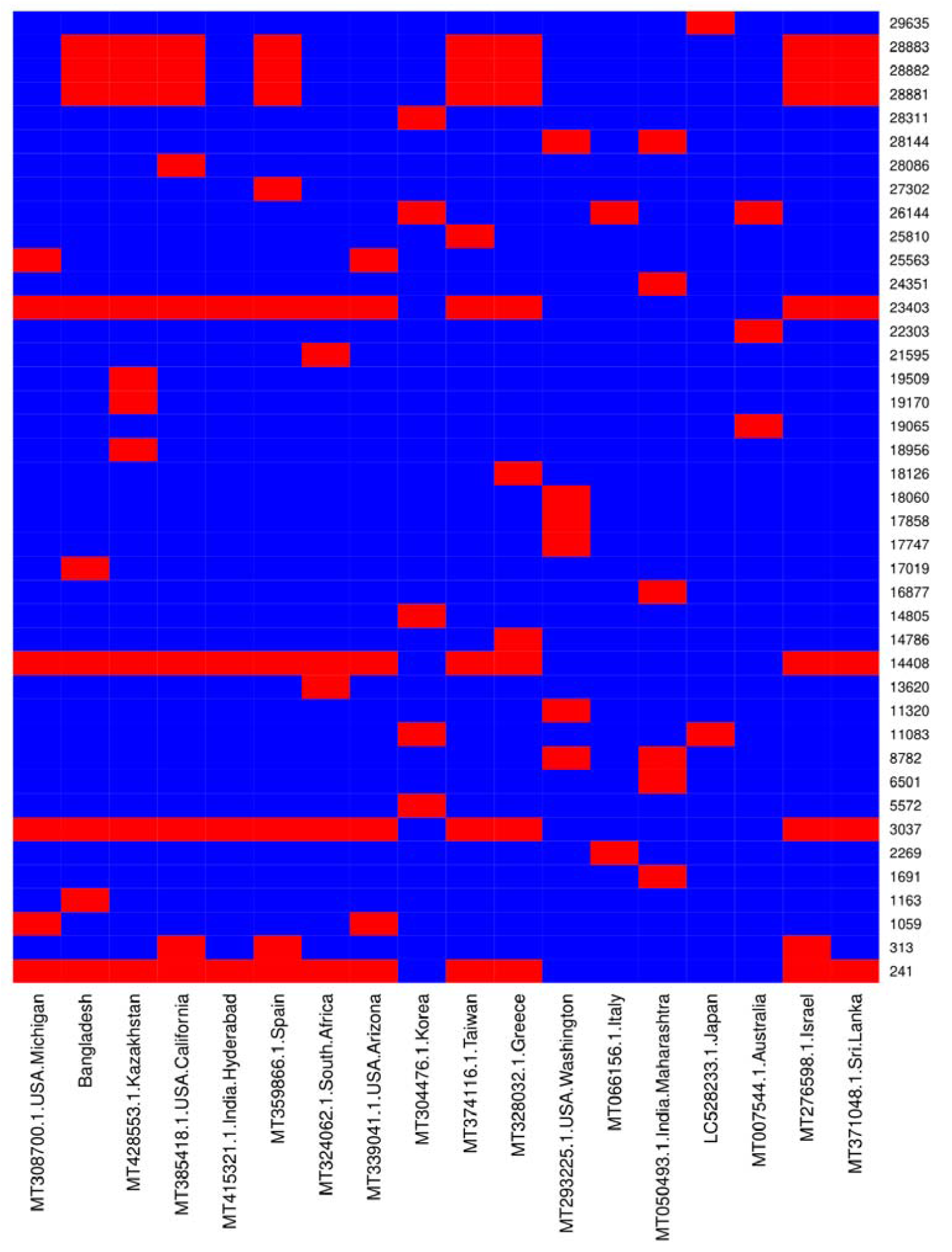
Binary heatmap showing all the variants identified using the BasebyBase tool. A total of 41 variants were identified in this way. The Bangladeshi isolate appears to share the most similarity with the isolate from Sri Lanka

**Figure 2A:**
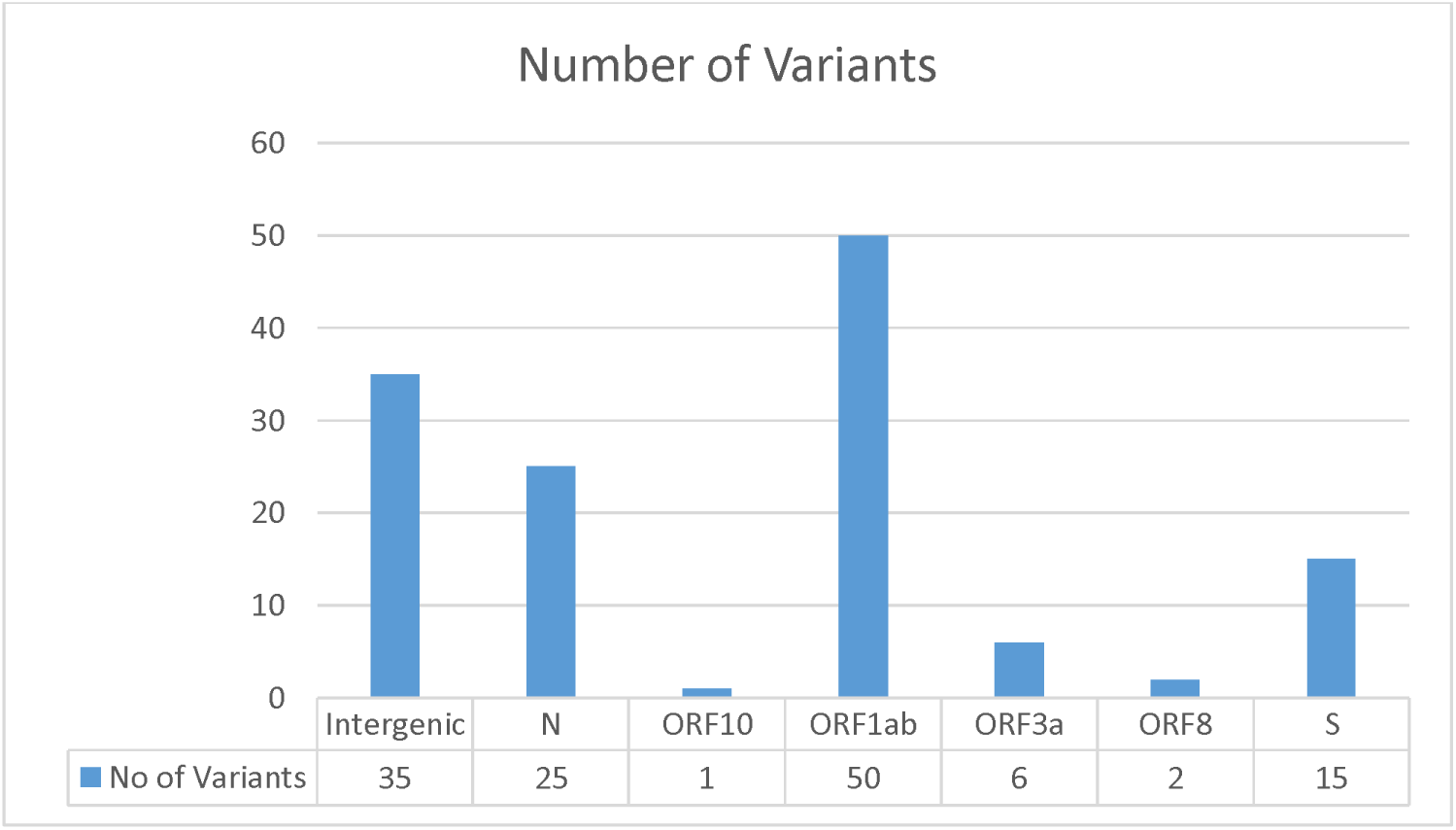
Figure 3: The number of variants per gene for all the isolates analyzed. ORF1ab had the most number of variants as should be expected because of its larger size. Interestingly the N protein had a comparatively high number of variants (22). This is somewhat unexpected for a structural protein for whom the organism should not be able to tolerate a high mutation rate.

**Figure 2B:**
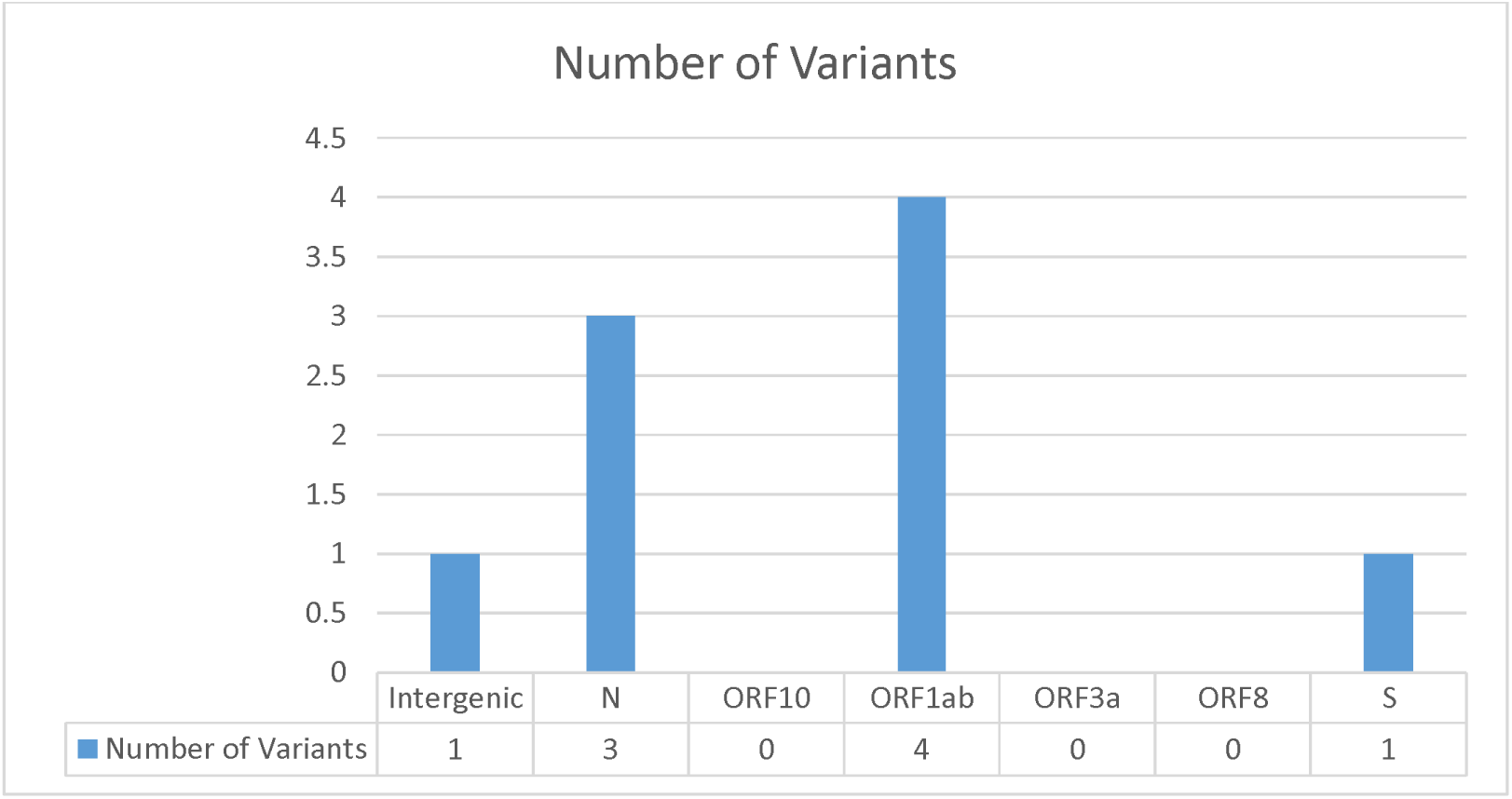
Number of variants per gene for the Bangladeshi SARS-CoV-2 isolate. As it can be seen, ORF1ab had the most number of variants, as can be expected because of its larger size.

The phylogenetic analysis revealed some interesting trends of its own. To begin with, the Banladeshi isolate was grouped in the same clade as isolates from Taiwan, Kazakhstan and Greece. This seems to reinforce what we saw with the BLAST results. The Taiwanese, Greek and Sri Lankan isolates occupied shorter branches than the Bangladeshi isolate, while the Kazakh isolate occupied a longer branch. It should be noted however the overall mean distance was 0 for this population of viruses, as can be deduced from the very small scale value. Despite the lack of apparent variation, we can still see some possible routes the virus may have taken when spreading across the globe and eventually reaching Bangladesh.

**Figure 4:**
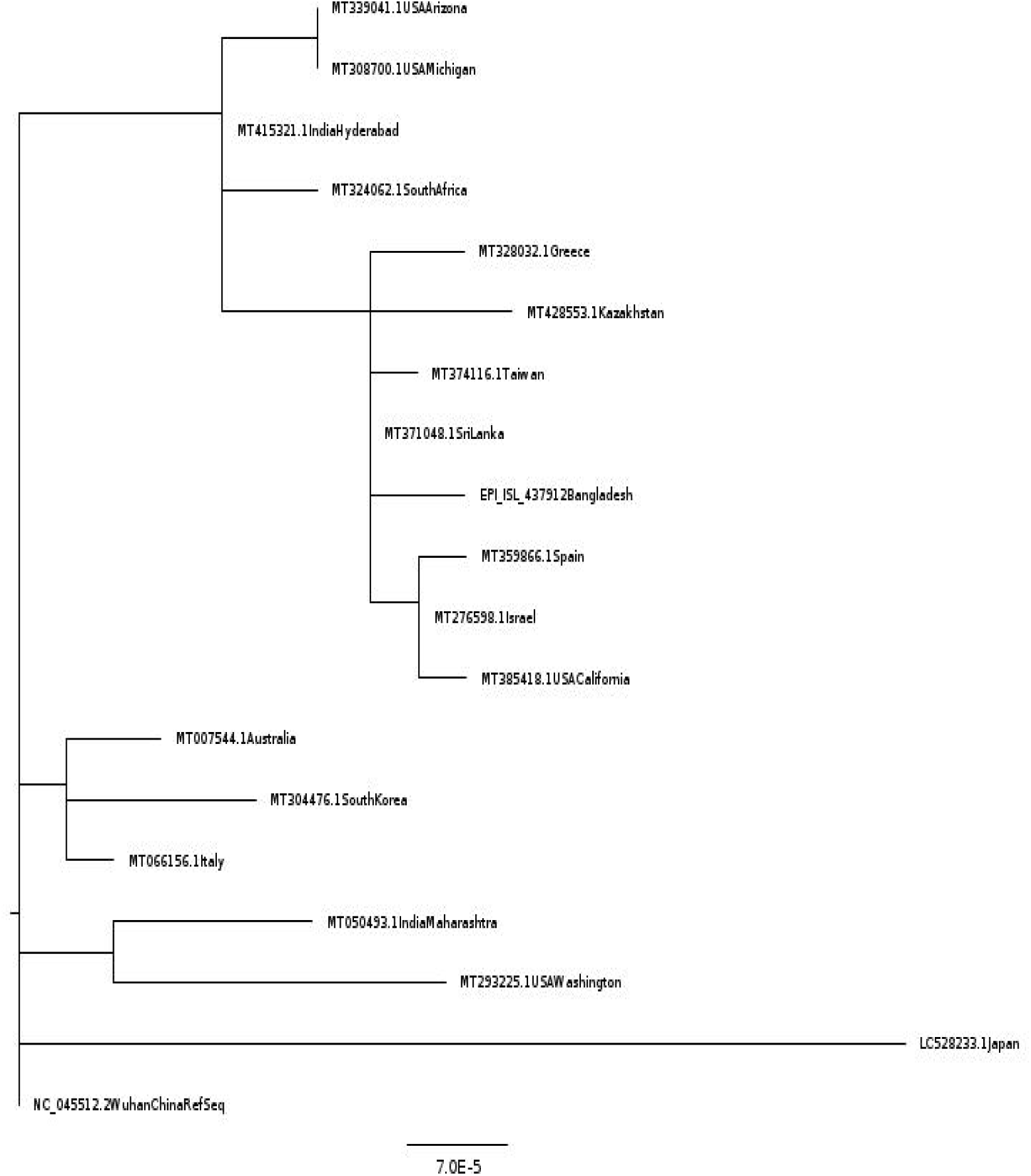
Phylogenetic tree compiled using sequence similarity between all our isolates. A very short scale value indicates these genomes are mostly identical barring a few nucleotides. The Bangladeshi isolate occupies one of the furthest nodes from the RefSeq, confirming it arose more recently.

If we observe the phylogenetic tree from the Wuhan RefSeq onwards, the first divergence event appears to give rise to the Japanese isolate, followed another that gives rise to three separate clades. One contains the Italian, South Korean and Australian isolates, another consists of the Indian (Maharashtra) and American (Washington) isolates and a third one that later gives rise to two more clusters. The first of these two clusters contains the isolates from Arizona and Michigan, while the second includes the Bangladeshi isolate. This observation that the Arizona and Michigan isolates and the Bangladesh isolate arise from the same divergence event and the fact that the two American isolates appear to be predate the Bangladeshi one, would seem to suggest that the virus first arrived in Bangladesh from USA. This seems to indicate that the initial speculation of the virus reaching Bangladesh through individuals arriving from Italy was perhaps incorrect.

## Discussion

Our goal behind this study was to try and understand the phylogenetic relationship between the Bangladeshi virus and those causing infections in other parts of the world, as well to characterize the unique genetic signature of this local strain.

The Bangladeshi appears to share a close sequence similarity with isolates from Taiwan, Kazakhstan and Greece, suggesting that the virus may have reached these countries from the same source; possibly Michigan or Arizona in the United States. The Sri Lankan isolate occupied the shortest branch in this cluster, possibly suggesting that it predates the Bangladeshi, Taiwanese, Greek and Kazakh isolates. Interestingly, the South African isolate, which shared almost complete sequence identity with the Bangladeshi isolate, appeared to predate it. This would indicate that perhaps the virus may have reached South Africa before Bangladesh. The Sri Lankan isolate makes for some interesting interpretation. Its shorter branch length points towards an earlier origin compared to the Bangladeshi virus. But the near 100% sequence identity would suggest otherwise. A plausible explanation may be that the virus both these countries from the source (most likely Michigan or Arizona), but it reached Sri Lanka before Bangladesh.

As for the SNPs, on average each isolate contained 6 SNPs. The Bangladeshi virus somewhat bucked the trend, containing 9 SNPs, tying with the Californian, Spanish and Greek viral genomes for the second highest among all the isolates analysed. The Kazakh viral genome had the highest number of variants at 10. The intriguing point to note here is that Kazakhstan and Greece have had comparatively fewer cases of infections (5207 and 2726 respectively). Spain and California have of course been much worse hit, but there we have to entertain the possibility of more than viral strain being active in those regions. The reason for that being both the Californian and Spanish isolate genomes we analysed were closely situated with the Bangladeshi isolate in the phylogeny. Given these countries reported their first infections much earlier than Bangladesh, it would seem unlikely that their viral isolates would appear to arise so late in the evolutionary course taken by this virus. This is backed up by the fact that this specific Spanish isolate genome was submitted on 17^th^ April and the Californian isolate genome on 23^rd^ April. It seems more likely that there may be multiple strains active in California and Spain and the ones we analysed arose much more recently. This is something that requires further analysis using more genomes from these countries. Hence it is wise to not correlate the statistics in those areas with the other countries mentioned. In summation, we believe the relationship between the number of mutations and the weakening of the virus is something that should be investigated. And our study provides at least some correlational evidence in its support.

The three N protein variants is another curious aspect of the Bangladeshi strain. It only appeared in the closely related isolates such as the Bangladeshi, Kazakh, Californian, Greek and Sri Lankan ones. The Israeli isolate also contained these. And the percentage of fatalities among infected patients is current a little over 1% for Israel; comparable with the Bangladeshi fatality percentage and considerably lower than the global. This invites the possibility of this, the three consecutive variants, possibly being a characteristic feature of a weaker strain of the virus. Again, the evidence right now is purely correlational. But this does still remain as another interesting future avenue to explore.

Lastly, the Spike protein variant at position 23403 was also found in the Bangladeshi virus. We are as of yet uncertain over what this could mean. Especially since contrary to previous claims, we also found this variant in other Asian isolates.

In conclusion, we believe the apparently higher rate of mutation in the Bangladeshi SARS-CoV-2 strain may hold possible clues pertaining to a mutation induced weakening of the virus. In addition we believe that the most likely route the virus used to get to Bangladesh was via either Michigan or Arizona in the USA. Sri Lanka also remains an outside possibility, but we believe the more likely scenario is that Sri Lanka and Bangladesh got the virus from the source, and that it arrived in Sri Lanka earlier than Bangladesh. Future studies should be putting more focus on how these variants impact the protein functions of the virus, so as to shed better light on how the pathogen is changing and what it means for our fight to bring the pandemic to an end.

## Supporting information

Supplemental Sequence File 1

## Supplementary Material

1. Supplementary Table 1: BLAST hits for Bangladeshi SARS-CoV-2 Isolate
2. Supplementary Table 2: BLAST results for multiple sequence alignment of all isolates used.
3. Supplementary File 1: MAFFT multiple sequence alignment results for all isolates.

